# Mutational analysis of the structure and function of the chaperone domain of UNC-45B

**DOI:** 10.1101/2020.01.03.894048

**Authors:** I. Gaziova, T. Moncrief, C. J. Christian, M. White, M. Villarreal, S. Powell, H. Qadota, G. M. Benian, A. F. Oberhauser

## Abstract

UNC-45B is a multidomain molecular chaperone that is essential for the proper folding and assembly of myosin into muscle thick filaments *in vivo*. We have previously demonstrated that its UCS domain is responsible for the chaperone-like properties of UNC-45B. In order to better understand the chaperoning function of the UCS domain we engineered mutations designed to: i) disrupt chaperone-client interactions by removing and altering the structure of the putative client-interacting loop and ii) disrupt chaperone-client interactions by changing highly conserved residues in the putative client-binding groove. We tested the effect of these mutations by using a novel combination of complementary biophysical (circular dichroism, intrinsic tryptophan fluorescence, chaperone activity, and SAXS) and *in vivo* tools (*C. elegans* sarcomere structure). Removing the client-holding loop had a pronounced effect on the secondary structure, thermal stability, solution conformation and chaperone function of the UCS domain. These results are consistent with previous *in vivo* findings that this mutation neither rescue the defect in *C. elegans* sarcomere organization nor bind to myosin. We found that mutating several conserved residues in the client-binding groove do not affect UCS domain secondary structure or structural stability but reduced its chaperoning activity. We found that these groove mutations also significantly altered the structure and organization of the worm sarcomeres. We also tested the effect of R805W, a mutation distant from the client-binding region. Our *in vivo* data show that, to our surprise, the R805W mutation appeared to have the most drastic effect on the structure and organization of the worm sarcomeres. In humans, the R805W mutation segregates with human congenital/infantile cataract, indicating a crucial role of R805 in UCS domain stability and/or client interaction. Hence, our experimental approach combining biophysical and biological tools facilitates the study of myosin/chaperone interactions in mechanistic detail.

**Statement of Significance:** The folding of myosin and the assembly of a functional sarcomere requires the chaperone UNC-45B. The molecular mechanism(s) for how UNC-45B assist in this assembly process or prevent stress-induced aggregation states are presently unknown. Answering this question is a problem at the core of muscle development and function. Here we developed a novel approach that combines biophysical and biological tools to study UNC-45B/myosin interactions in mechanistic detail. Our approach may provide critical insights into the molecular nature of the pathogenesis of many muscle disorders stemming from mutations in sarcomeric proteins including skeletal myopathies and cardiomyopathies, and possibly the age-associated decline in muscle mass and function found in the elderly known as sarcopenia.

## 1. INTRODUCTION

The arrangement of not just the thick and thin filaments, but numerous other proteins into the exacting arrangement of the semi-crystalline lattice making up the sarcomere is essential for muscle contractile function. This process is partially autonomous, an intrinsic property of its component proteins. However, the assembly of a functional sarcomere requires molecular chaperones (1). These serve to prevent aggregation of unfolded intermediates and according to recent evidence, to help assemble the sarcomere (2–4). The molecular mechanism(s) for how chaperones assist in this assembly process or prevent stress-induced aggregation states are presently unknown. Answering this question is a problem at the core of muscle development and function (5, 6). This understanding may provide critical insights into the molecular nature of the pathogenesis of many muscle disorders stemming from mutations in sarcomeric proteins including skeletal myopathies and cardiomyopathies, and possibly the age-associated decline in muscle mass and function found in the elderly known as sarcopenia. Amongst the sarcomeric chaperone proteins known to be involved in the folding of myosin is UNC-45 (2, 3, 7–10). Although *C. elegans* and Drosophila each have only one unc-45 gene, vertebrates have two unc-45 genes, one (UNC-45B), expressed only in striated muscle cells, and the other (UNC-45A) expressed in multiple cell types (8).

UNC-45B is a UCS-domain containing protein that functions as a myosin chaperone promoting the folding of the myosin head (or S1 fragment) as well as coordinating the assembly of thick filaments during muscle development (2, 4, 7–11).

Our previous data demonstrates that UNC-45B is a unique multidomain chaperone where each domain has a clearly defined and distinct function (**Fig. 1**): i) the TPR domain binds specifically to Hsp90 (11–13); ii) the central domain, binds the myosin head and/or neck (14)) and inhibits the actin translocation function of myosin (15); iii) the conserved UCS domain is alone capable of preventing the aggregation of its client protein, the myosin head, and it is responsible for the chaperone-like properties of UNC-45B (14).

**Figure 1.**
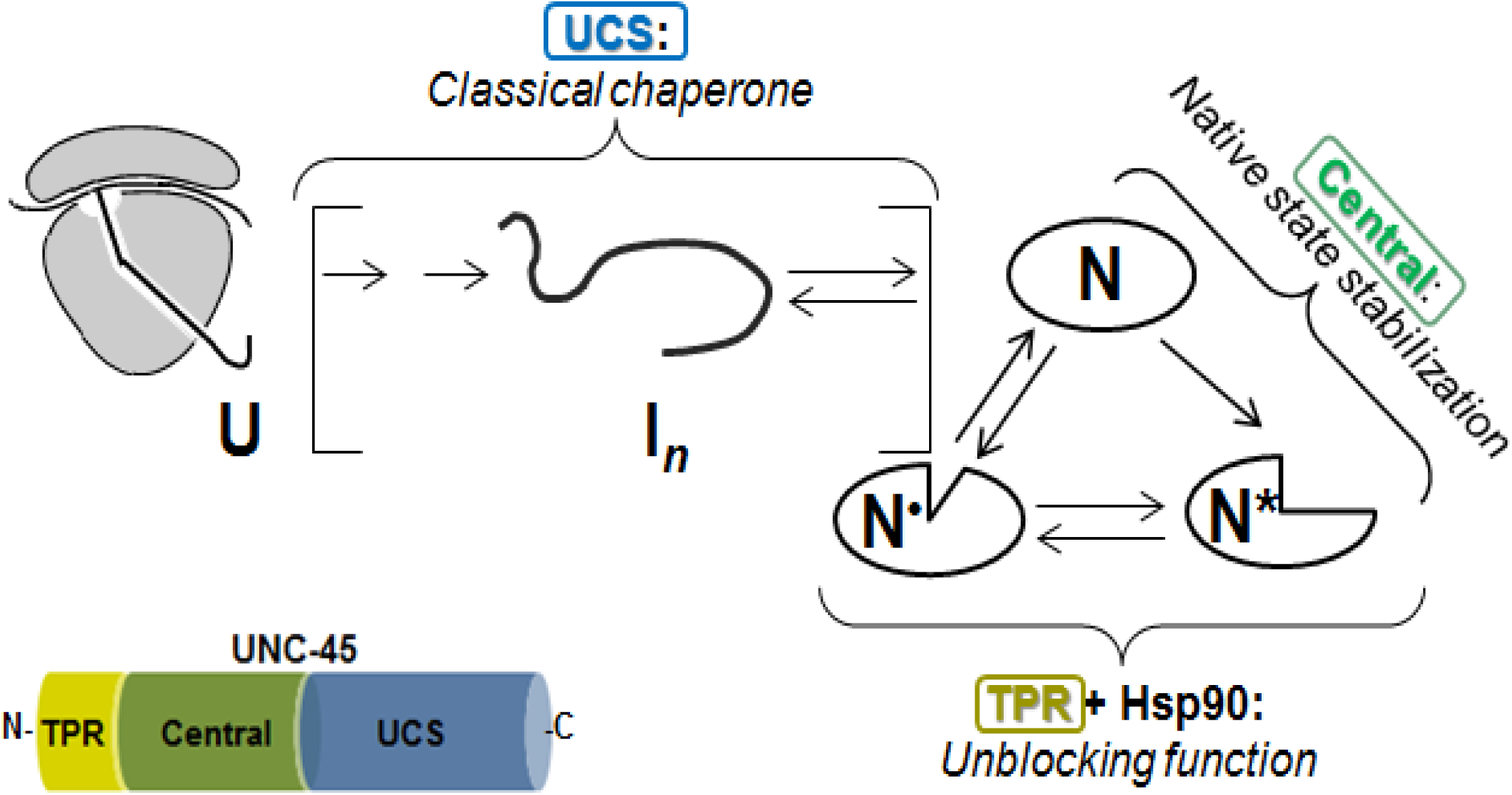
**Proposed model for the function of UNC-45 domains in myosin interactions.** The **UCS domain** (in blue) prevents misfolding and aggregation of unfolded (U) and partially folded intermediates (I*_n_*) of ribosome (in gray) nascent chains. The **central domain** (in green) stabilizes one of several native states (N*), incapable of translocating actin (diagrammatic only, many more native states of myosin are known). The **TPR domain** (in yellow) recruits the molecular chaperone Hsp90 to unblock the action of UNC-45, allowing resumption of actin gliding.

Structurally, the myosin-interacting element shared by UCS proteins consists of nine copies of the armadillo domain (3, 8, 16–20). Although no human UNC-45 structure is available, homology models have been created via *in silico* molecular modeling and simulations (21). **Fig. 2A** shows a homology model of the human UCS domain of UNC-45B depicting armadillo repeats 9 to 17 (armadillo repeats 1-8 reside in the central domain). These armadillo repeats consist of 3 alpha helices (a short H1 of roughly two helical turns followed by two longer H2 and H3 helices) with an average of 42 residues with diverse sequences, but highly conserved structures (**Fig. 2B**). These helices arrange in a superhelical structure defining a protein-binding groove that is conserved in many armadillo repeat containing proteins (17, 22–24). In human UNC-45B this groove comprises highly conserved residues in the UCS domain, such as the NYE…LTNL and DRLK sequences in repeats 13 and 15, respectively (highlighted in green in **Fig. 2B**). When mapped onto the human UCS homology model these highly conserved amino acid residues in helix 13H3 form a large surface patch in the groove (highlighted in red in **Fig. 3A**). Based on the marked homology of the UCS domain to β-catenin, Gazda et al. (2013) (17) predicted that this groove may serve as a myosin-binding surface and were able to identify key residues for myosin interaction. By mapping a β-catenin ligand onto the *C. elegans* UNC-45 UCS domain they predicted that N758 (N745 in human sequence) should be critical for tethering the main-chain of the captured substrate, whereas the nearby Tyr750 (Y737 in human) should contribute to the hydrophobic character of the outer rim of the UCS groove (**Fig. 3).** *In vivo* mutagenesis analysis revealed that in contrast to a Y750W mutant, which is still capable to restore muscle function, a N758Y mutant failed to rescue a temperature-sensitive *unc-45* mutant (in worms carrying the ts allele *unc-45(m94, mutation E781K*) grown at the nonpermissive temperature. Further analysis of *C. elegans* mutations affecting myosin-binding groove by the same group confirmed that UNC-45 carrying Y750W mutation was able to support production of soluble myosin motor domain when coexpressed in insect cells, even though to less extent than wt UNC-45 (20). N758Y unfortunately failed to express in insect cells efficiently, suggesting structure destabilizing nature of this mutation.

**Figure 2.**
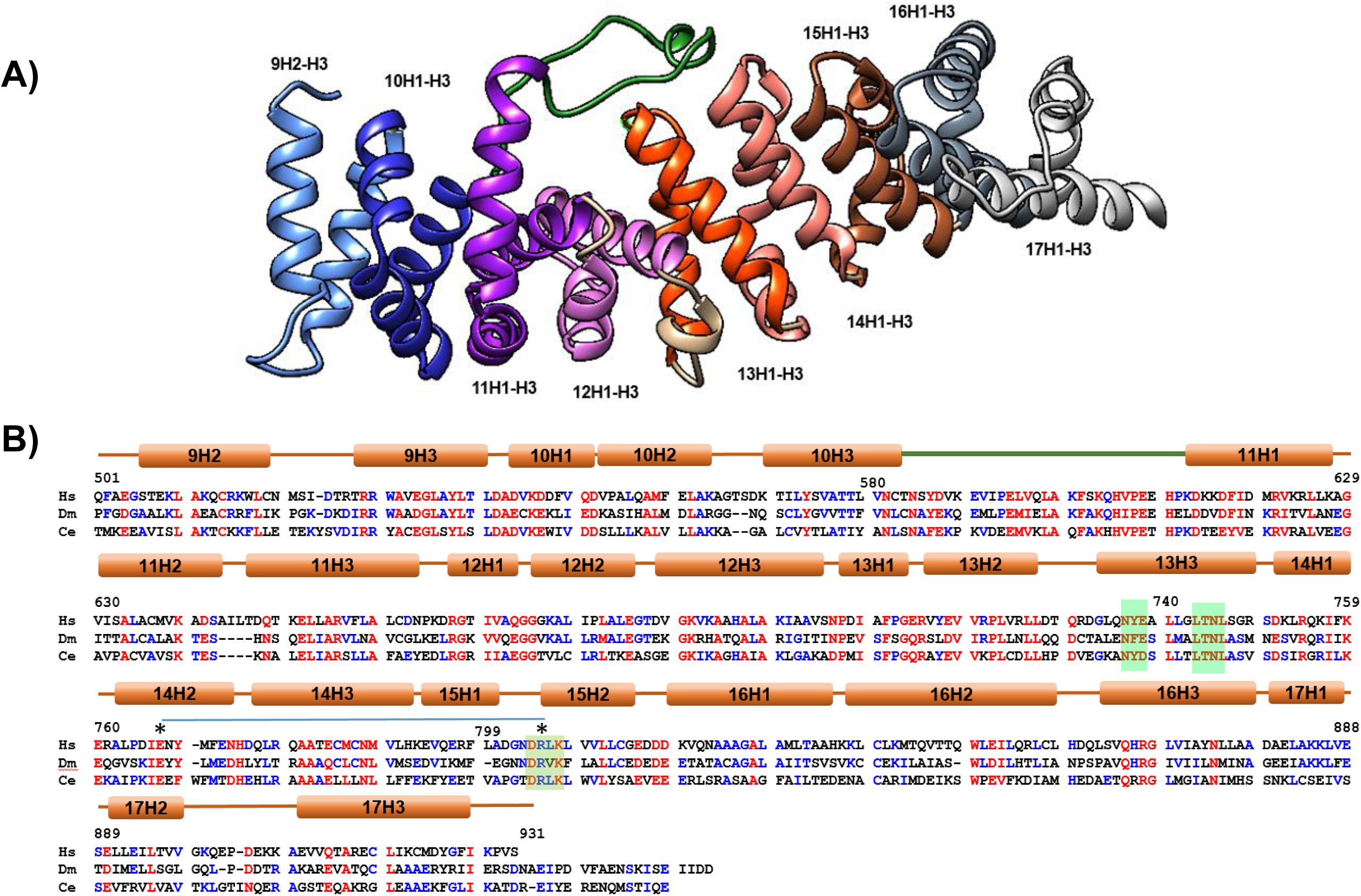
**A)** Homology model of the human UCS domain based on the *C. elegans* 3D structure (PDB ID: <u>4i2z</u>), highlighting the different ARM repeats (9 to 17). Phyre2 ((58), SWISS-MODEL (59) and Chimera (60) were used for making the human model. **B)** Sequence alignment of the UCS domains from UNC-45 proteins from Homo sapiens (Hs; E1P642), Drosophila melanogaster (Dm; Q9VHW4) and *C. elegans* (Ce; Q09981). Red residues are highly conserved in all three sequences; blue are low consensus residues. Green boxed residues are highly conserved residues in the groove region. **\*** denotes conserved inter-repeat interacting ion pair Glu768 and Arg805 residues. The overall sequence identity between the HsUNC-45 and the other listed UNC-45 proteins is about 36%. ClustalOmega (61) and Multalin (62) were used for the alignment.

**Figure 3.**
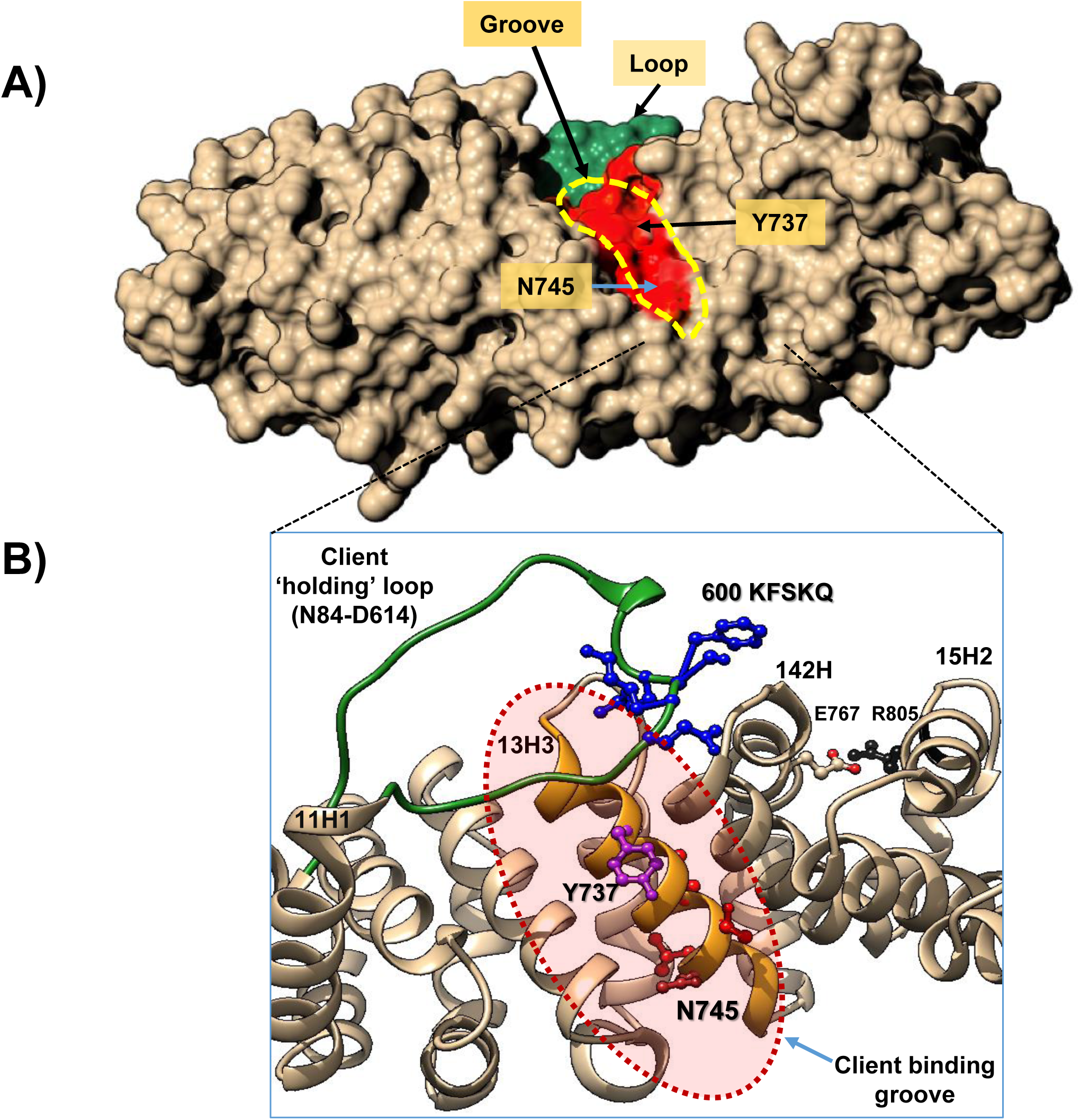
**A)** Surface representation of the human UCS domain highlighting the loop region (in green) and the highly conserved residues in the groove region, 736 NYEaLTNL 746) (red patch). **B)** Location of the different residues mutated in the UCS domain: deleted loop in green (N584-D614), mutations in the loop 600 KFSKQ 604 in blue, Y737 (in purple), N745 (in red) and R805 (in black). Chimera (60) was used to make the representation.

Another potential important myosin-interacting region of UCS proteins is a flexible loop with minimal electron density conserved in the UNC-45 structures of *C. elegans* and *D. melanogaster,* which has been shown to be susceptible to trypsin digestion in the Drosophila protein (19). This flexible loop is colored green in **Figs. 2** and **3**, and when deleted should disrupt the substrate-binding cleft and abrogate the interaction with myosin and it was found that a Δ602–630 mutant (N584-D614 in human sequence) neither rescue the defect in sarcomere organization nor bind to myosin in immunoprecipitation experiments (17). These data indicate that the loop may sense and capture an unfolded portion of myosin, and act as a “trap door” to capture exposed myosin peptides within the UCS groove (8). We propose that the groove is the inside of a box and the loop is the lid of the box.

Another suitable candidate for UCS domain destabilization and reduction of its ability to bind myosin head is Arg805 since based on x-ray crystal structure of Drosophila UNC45, R803 (R805 in human) has been implicated to form inter-repeat ion pair interaction with Dm E766 (E767 in human) and thus stabilize ARM repeats 18 and 19 (repeats 14 and 15 in human UNC-45B) (19). Consistent with this idea, autosomal dominant negative mutation R805W in human UNC45B has been associated with congenital/infantile cataract in three generations of a Danish family (25).

In this work we used mutagenesis to test the following hypotheses: i) that the putative client-interacting loop in UCS functions as a sensor and holder for the unfolded or partially folded client polypeptide chain (‘client-holding loop’) and ii) that the putative ‘client-binding groove’ region in UCS functions as a binding site for the client polypeptide chain. We used a combination of complementary assays that probe the effect of mutations on: i) its structure and stability by Circular Dichroism (CD); ii) its conformational changes (by using tryptophan fluorescence); iii) its client chaperoning activity (by using heat-inactivation protection assays); iv) its overall shape and conformation in solution (by using Small angle X-ray scattering (SAXS)); and v) assessing the *in vivo* effects in *C. elegans*.

## 2. MATERIALS AND METHODS

### Mutagenesis, Protein Expression and Purification of proteins

The UCS domain constructs (wild-type and mutants) of UNC-45B (residues 501-931) from *Homo sapiens* were codon optimized for expression in *Escherichia coli*, synthesized (GenScript, Piscataway, NJ) and subcloned into a pET-28 vector (EMD Millipore, Billerica, MA) or pET30A (Novagen) respectively. The N745W, Y737W and R805W substitution in were introduced into UCS-wt using Q5® Site-Directed Mutagenesis Kit (NEB). Recombinant protein expression was induced in BL21 DE3 when optical density (OD600) reached 1-1.2 with 0.02 mM IPTG for 16 h at 15 °C. Harvested cells were resuspended in lysis buffer (50 mM Tris-HCl, 50 mM NaCl, 40 mM imidazole, 2 mM TCEP, 10% glycerol, pH 8.0) containing 1 mg/ml lysozyme and incubated for 30 min at RT. To promote lysis cells further samples were sonicated 6 x 20 s with 20 s breaks (200 – 300 W) on ice. To break down DNA and reduce viscosity, 1 µl of Universal nuclease (Pierce; 125U) and 10 of DNaseI (NEB; 20U) were added to the sample and incubate for 10 min at RT. Insoluble material was removed by centrifugation at 30,000 x g for 30 min at 4 °C and supernatant was filtered through 0.45 µm syringe filter afterwards. Supernatant was incubated with lysis buffer equilibrated HisPur Ni-NTA Resin (Thermo Scientific) for 1 hour at 4°C. Resin with bound proteins were collected by centrifugation at low force (700 x g for 2-3 min), washed by 4 washes with ice cold wash buffer (50 mM Tris-HCl, 50 mM NaCl, 60 mM imidazole, 2 mM TCEP, 10% glycerol, pH 8.0) and with last wash applied to the column cartridge. His-tagged recombinant protein was eluted in 1 ml fractions with elution buffer (50 mM Tris-HCl, 50 mM NaCl, 250 mM imidazole, 2 mM TCEP, 10% glycerol, pH 8.0) and immediately 4 µl of EDTA pH 8.0 were added. Eluted fractions containing significant portion of recombinant protein were pooled and dialyzed against storage buffer (50 mM Tris-HCl, 50 mM NaCl, 2 mM EDTA, 2mM TCEP, 10% glycerol, pH 8.0) overnight at 4°C. Proteins were concentrated to 20-40 µM, flash frozen and long term stored at −80°C. For SAXS analyses, freshly prepared recombinant proteins were concentrated to 1 mg/ml and any aggregates were removed by ultracentrifugation at 100,000 x g for 30 min at 4°C.

### Circular Dichroism

The far UV CD spectra of the UCS constructs were recorded on a Jasco J-815 Spectrometer. The protein concentration was 1 μM in 30 mM phosphate buffer pH 7.4, 100 mM KCl, 1 mM MgCl_2_, 1mM TCEP buffer. A 0.1 cm path length cuvette was used. The temperature was automatically adjusted in 5 °C steps from 20 to 60 °C with 25 min of equilibration at each point after reaching constant temperature. The fraction of unfolded protein was calculated, using the following equation:

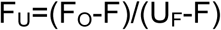

where F_U_ is fraction unfolded, F_o_ is the value of ellipticity (θ) at 222, F is the average of first value or average of few values of ellipticity where protein is folded, U_F_ is the average of last value or average of last few values of ellipticity where protein is unfolded. The melting temperature (Tm) was estimated by fitting the value in the Boltzmann sigmoid equation, which assumes a two state model (26, 27).

### Intrinsic fluorescence

1 μM protein in 30 mM phosphate buffer pH 7.4, 100 mM KCl, 1 mM MgCl_2_, 1mM TCEP buffer was placed in a quartz cuvette in a Fluorolog-3 fluorescence spectrometer (Horiba Jobin Yvon, Kyoto, Japan) equipped with a 40 W temperature controller (Wavelength Electronics Bozeman, MT). The protein was excited at 285 nm (tryptophan absorption) and the emission spectra from 300–600nm. The temperature was adjusted in 5 °C steps from 20 to 60 °C with 25 min of equilibration at each point after reaching constant temperature. At the end of each experiment, the sample was cooled back to 20°C to check for recovery of the original signal. In order to estimate the melting temperature (Tm), the intensity of fluorescence at its maximum wavelength (λ_max_) was plotted as a function of the temperature. Background spectra from only the buffer were recorded and subtracted at each temperature.

### Small angle X-ray scattering (SAXS)

All SAXS data were collected using a Rigaku BioSAXS-1000 camera and ASC-96 Automatic Sample Changer on a FR-E++ Cu x-ray source (Rigaku Americas Corporation, Woodlands, TX). Each protein sample was buffer exchanged and concentrated in 50 mM Tris-HCl, 50 mM NaCl, 2 mM EDTA, 2mM TCEP, 10% glycerol, pH 8.0. The concentrator flow-though was used as the matching buffer, after thorough washing of the concentrator. A dilution series of 4.0, 2.0, and 1.0 mg/ml for the UCS WT protein and a dilution series of 1.0, 0.67, and 0.50 mg/ml for the UCS DEL protein, each with a matching buffer, was collected. At each concentration, 70 μl of sample plus a matching buffer were manually pipetted into separate tubes of an eight-tube PCR strip, sealed, and loaded in to the ASC-96 sample changer. The samples in the ASC-96 were held at the same temperature as the sample capillary cell, 10° C. For each sample a series of 30 minute exposures were collected, and averaged in SAXLab to produce separate sample and buffer curves of from 6 to 12 hours total exposure (**Table S1**). Data were collected in the range 0.014 Å^-1^ < q < 0.68 Å^-1^ for WT and 0.011 Å^-1^ < q < 0.68 Å^-1^ for DEL, and analysis used all significant data to 0.50 Å-1. The 4 mg/ml WT data were radiation sensitive and only the first 6 hours were averaged. The other samples did not display any radiation-induced or time-dependent changes. Buffer subtraction, absorption correction, and MW calibration were performed using the SAXNS-ES server (http://xray.utmb.edu/SAXNS), which also uses an concentration and intensity independent method to determine the MW (28). Data analysis, including merging of curves, was performed with the Primus program and the P(r) was calculated using DATGNOM, both from the ATSAS suite (29). The *ab initio* molecular shape was generated from an average of 15 DAMMIF (30) runs, using the saxns_dammif utility or an average of 4 clustered GASBOR (31) models, using the saxns_gasbor utility, damclust, and damaver. Both P1 and P2 symmetry ab initio models were created. Rigid body modeling of the dimer was performed using CORAL (29) with the 4i2z PDB model. The concentration-dependent mixture analysis was performed using Oligomer (Konarev, 2003) with the 4 mg/ml WT Dimer model or the 0.50 mg/ml DEL Dimer model from CORAL. The monomer-dimer equilibrium was modeled using SASREFMX and the dilution series data. The analysis optimizes the dimer configuration, based on the provided monomer structure, while adjusting the concentration-dependent monomer/dimer fractions to minimize the total χ^2^ deviations of all three data sets simultaneously.

### S1 Mg-ATPase heat inactivation protection assay

0.4 µM Myosin Motor protein S1 Fragment (Rabbit Skeletal Muscle, Cytoskeleton #CS-MYS04) was incubated either alone or with 2 µM UCS proteins for 10 minutes either at 42°C or on ice in Mg-buffer (10mM Hepes-KOH, 100mM KCl, 10 MgCl_2_, 0.3 mM EGTA, 2mM TCEP, pH 7.4). Heat exposed samples were cooled down on ice for 1 minute and mixed 1:1 with 1 mM ATP in Mg-buffer to initiate ATP hydrolysis for 15 min at room temperature. S1 ATPase activity was quenched with PiColorLock™ Gold Reagent (Expedeon Inc., San Diego CA 92121) and free Pi detection was carried out according to the manufacturer’s recommendation. Samples were read at wavelength 620 nm using a FLUOstar Optima Microplate reader (BMG Labtech Inc.). The assay was carried in triplicates, obtained readings were blank corrected and used to calculate relative activity to S1.

### Citrate Synthase Activity Assay

Citrate Synthase (CS) Activity Assay was adapted from Buchner at al, (32, 33). CS catalyzes the reaction of oxaloacetic acid (OAA) and acetyl coenzyme A (Acetyl-CoA) to citrate and CoA. The enzyme was determined by a colorimetric test using DTNB, which reacts with the free thiol groups of the reaction product CoA. This reaction was followed in a spectrophotometer at 412 nm. An ammonium sulfate suspension of porcine citrate synthase (Sigma; C3260) was dialyzed against 50 mM Tris-HCl, 2 mM EDTA, 10% glycerol, pH 8.0 overnight at 4°C, concentrated to about 100-200 µM using Amicon Ultra centrifugal filters with 30 kDa cut-off limit, and stored at −80°C.

Prior to activity measurement, CS was diluted to 0.2 µM in 40 mM Hepes-KOH, pH 7.4 and 2-4 µl of the enzyme was mixed with 100 µl TE buffer containing 100 nM Ellman’s Reagent (DTNB) (Invitrogen) and 150 nM Acetyl coenzyme A (Sigma) in a 96 well plate. To initiate the reaction, freshly prepared oxaloacetic acid, pH ∼7.4 was diluted to 100 nM in 20 mM Tris HCl, 100mM KCl and 100 µl of the mixture was added to the reaction. Immediately, absorbance at 412nm was measured at RT every 30s for 12 cycles using a FLUOstar optima Microplate reader (BMG Labtech Inc.). After subtracting blank readings (reaction mixture without CS) from obtained values, change in CS activity was determined from the linear range of the initial absorbance increase per 1 minute at two different time points (60-0s and 90s-30s).

### Influence of UCS Mutants on the Thermal Inactivation of Citrate Synthase

UNC45B, UCS and its mutant variants were mixed with CS and diluted in 10:1 ratio in 20 mM Tris HCl, 100mM KCl, pH 8.0 (at a concentration of 2 µM and 0.2 µM, respectively. Each mix, of CS and chaperones, as well sample of CS alone were split into 5 different tubes, which were exposed to 42 °C heat shock for 0, 5, 10, 15 and 20 minutes (**Figure 7**) or incubated for 10 min at 42°C (**Figure 8)**. Upon heat shock, samples were cooled down to 4°C, briefly centrifuged and CS activity was performed as described in the previous methods section. Assays were run in triplicates in at least two independent experiments.

### Plasmid construction for *C. elegans*

A full-length cDNA for wildtype *unc-45* was amplified by PCR (**Table S2**) in three fragments from the wild type cDNA library RB2 (kindly provided by Robert Barstead). The fragments were ligated into a pBluescript(pBS) vector using either Sma I and Sal I (fragments #1 and #2) or just Sal I (fragment #3) and verified by sequencing. Fragments #1 and #2 were ligated together with pBS after Sal I and Nae I digestion. Fragment 1+2 was then digested out of pBS using Sma I and Sal I and ligated into the pKS-HA vector (34) digested with Eco RV and Sal I. Fragment #3 was then digested out of pBS with Sal I and ligated into 3XHA-pBS-fragment 1+2, also digested with Sal I. 3XHA-*unc-45* fragments1+2+3 was then digested out of pBS with Spe I and ligated into pPD95.86 (a gift from A. Fire, Stanford University, Stanford, CA), which contains the *myo-3* promoter, digested by Nhe I. Mutant *unc-45* UCS fragments were created and amplified from the pBS-fragment #3 (UCS) plasmid using primers with the intended mutation in combination with the original fragment #3 primers. The UCS fragments containing the correct mutations were then ligated into the 3XHA-pBS vector containing the rest of the *unc-45* sequence (described above). The four mutant 3XHA-*unc-45* sequences were then cut out of pBS with Spe I and ligated into pPD95.86 digested by Nhe I.

### *C. elegans* Strains

Standard growth conditions for *C. elegans* were used (35). For wild type and each of the mutant *unc-45* plasmids, three independently-generated extrachromosomal array transgenic lines were created by SunyBiotech Corporation by co-injection into wild type N2 Bristol of pPD95.86-HA-UNC-45 and the transformation marker plasmid pTG96 (psur-5::sur-5::nlsGFP, that expresses GFP in most somatic cell nuclei). These transgenic lines have the following strain names: For UNC-45 wild type, GB297-299; for UNC-45 Groove, GB300-302; for UNC-45 R805W, GB303-305; for UNC-45 Pro, GB306-308; and for UNC-45 2xW, GB309-311.

### Immunostaining in adult body-wall muscle

Adult nematodes were fixed and immunostained according to the method described by Nonet et al. (36) and described in further detail by Wilson et al. (37). The following primary antibodies were used: anti–MHC A at 1:200 (mouse monoclonal 5-6 (38)), anti–UNC-95 at 1:100 (rabbit polyclonal Benian-13 (34), anti-GFP at 1:200 (rabbit polyclonal (Invitrogen, Thermo Fisher Scientific A-11122)), and anti-HA (mouse monoclonal clone HA-7 (cat. no. H3663, Sigma Life Science)). Secondary antibodies, used at 1:200 dilution, included anti-rabbit Alexa 488 (Invitrogen, Thermo Fisher Scientific) and anti-mouse Alexa 594 (Invitrogen). Images were captured at room temperature with a Zeiss confocal system (LSM510) equipped with an Axiovert 100M microscope and an Apochromat ×63/1.4 numerical aperture oil immersion objective in ×2.5 zoom mode. The color balances of the images were adjusted by using Photoshop (Adobe, San Jose, CA).

### Statistical analyses

Unless otherwise stated, data are reported as mean ± standard error of the mean. Student’s paired t-test (2-tailed) was applied to determine the significance of the differences. Statistical significance was assigned as not-significant for P > 0.05 and * for P ≤ 0.05.

## 3. RESULTS

### UCS domain engineered mutations and predicted effects

Guided by the data presented in the Introduction section we have mapped putative key amino acid regions in the UCS domain that may mediate its myosin S1 binding and chaperoning functions. Some of the predicted effects of the mutations are shown in **Table I**. For example, we hypothesize that deletion of the ‘myosin-holding loop’ (a highly disordered region, aa N584-D613, “DEL” mutant) should disrupt the interaction with the myosin head. We hypothesize that increasing the rigidity of the loop by adding prolines (“PRO” mutant) in a partially conserved sequence (KFSKQ) should affect the chaperoning activity of the UCS. Proline-rich sequences are known to form rigid and extended structures that are fairly resistant to deformation or rotation (39) (40, 41); hence a more rigid loop should make the lid of the box close slower, resulting in a reduced chaperone activity.

**Table I.**
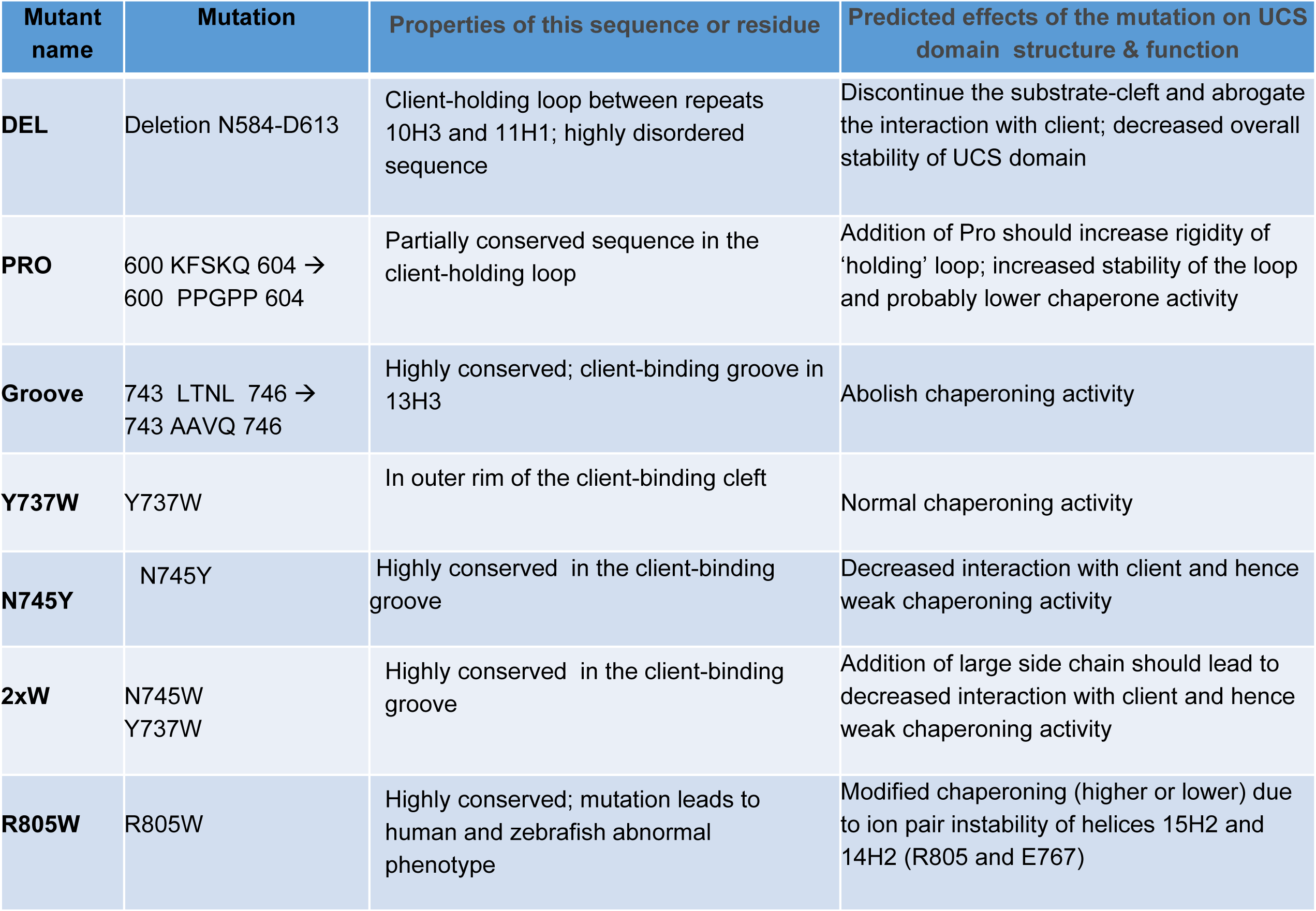

| Mutant name | Mutation | Properties of this sequence or residue | Predicted effects of the mutation on UCS domain structure & function |
| --- | --- | --- | --- |
| **DEL** | Deletion N584-D613 | Client-holding loop between repeats 10H3 and 11H1; highly disordered sequence | Discontinue the substrate-cleft and abrogate the interaction with client; decreased overall stability of UCS domain |
| **PRO** | 600 KFSKQ 604 → 600 PPGPP 604 | Partially conserved sequence in the client-holding loop | Addition of Pro should increase rigidity of 'holding' loop; increased stability of the loop and probably lower chaperone activity |
| **Groove** | 743 LTNL 746 → 743 AAVQ 746 | Highly conserved; client-binding groove in 13H3 | Abolish chaperoning activity |
| **Y737W** | Y737W | In outer rim of the client-binding cleft | Normal chaperoning activity |
| **N745Y** | N745Y | Highly conserved in the client-binding groove | Decreased interaction with client and hence weak chaperoning activity |
| **2xW** | N745W<br>Y737W | Highly conserved in the client-binding groove | Addition of large side chain should lead to decreased interaction with client and hence weak chaperoning activity |
| **R805W** | R805W | Highly conserved; mutation leads to human and zebrafish abnormal phenotype | Modified chaperoning (higher or lower) due to ion pair instability of helices 15H2 and 14H2 (R805 and E767) |

**Table II.**

| **Protein Name** | **% alpha, beta Structure** | **Stability (T<sub>m</sub>, °C)**<br>**CD; Trp** | **S1 assay**<br>**% Protection**<br>**(normalized)** | **CS assay**<br>**(%Protection**<br>**(normalized)** | **Worm**<br>**Sarcomere**<br>**phenotype** |
| --- | --- | --- | --- | --- | --- |
| **WT** | 66, 6 | 40; 27 | 100 | 100 | normal |
| **DEL** | 31, 36 | 43; 36 | 39 | 63 | - |
| **PRO** | 69, 6 | 40; 30 | 40 | 100 | Disrupted<br>(2/3) |
| **Groove** | - (not soluble<br>in *E. coli*) | - | - | - | Disrupted<br>(2/3) |
| **Y737W** | 68, 5 | 40; 26 | 51 | 86 | - |
| **2xW** | 69, 6 | 40; 33 | 61 | 73 | Disrupted<br>(2/3) |
| **R805W** | 70, 6 | 40; 33 | 47 | 77 | Disrupted<br>(3/3) |

We also hypothesize that mutations of key conserved residues in the putative ‘myosin-binding groove’ should reduce the chaperone activity of UCS. Residues Y737 and N745 (**Fig. 3**) are predicted to play an important role in the interaction of the unfolded or partially unfolded client protein with the groove (17); hence, the bulky side chains of the double mutant Y737W/N745W (“2xW”) should lead to a decreased interaction with client and also a weak chaperoning activity. We also mutated Arg 805 to Trp to test the idea that the stability of the salt bridge between R805-E767 (**Fig. 2B**) plays an important role in UCS-client interactions, as suggested by Hansen et al. (25).

In order the test these hypotheses we made recombinant UCS constructs with the mutations listed in **Table I**. We found that all proteins but the “Groove” construct expressed well as soluble proteins in *E. coli*. The “Groove” construct expressed as an insoluble protein in inclusion bodies, indicating that this mutation leads to misfolding.

### Effect of mutations on secondary structure and stability

In order to determine whether the mutant proteins folded to similar conformations to the WT we collected circular dichroism (CD) spectra in the far-UV region (200–250 nm). The far-UV CD spectrum obtained at 20 °C for the WT UCS domain (**Fig. 4**) is typical for alpha-helical proteins, showing minima at 208 and 222 nm, which is consistent with the available crystal structure data that show that it is composed almost entirely of armadillo repeats (17–19). As seen in **Fig. 4** the mean residual ellipticity traces of the PRO, Y737W, 2xW and R805W mutant proteins are very similar to the WT and secondary structure fitting using the K2D3 webserver suggests very little perturbation of the strong alpha-helical structural component (about 68% on average). In contrast, and to our surprise, the spectrum for the DEL mutant significantly deviates from the wild type, exhibiting a decrease in alpha helix character (to 31%) and increase in beta sheet character (to 36%).

**Figure 4.**
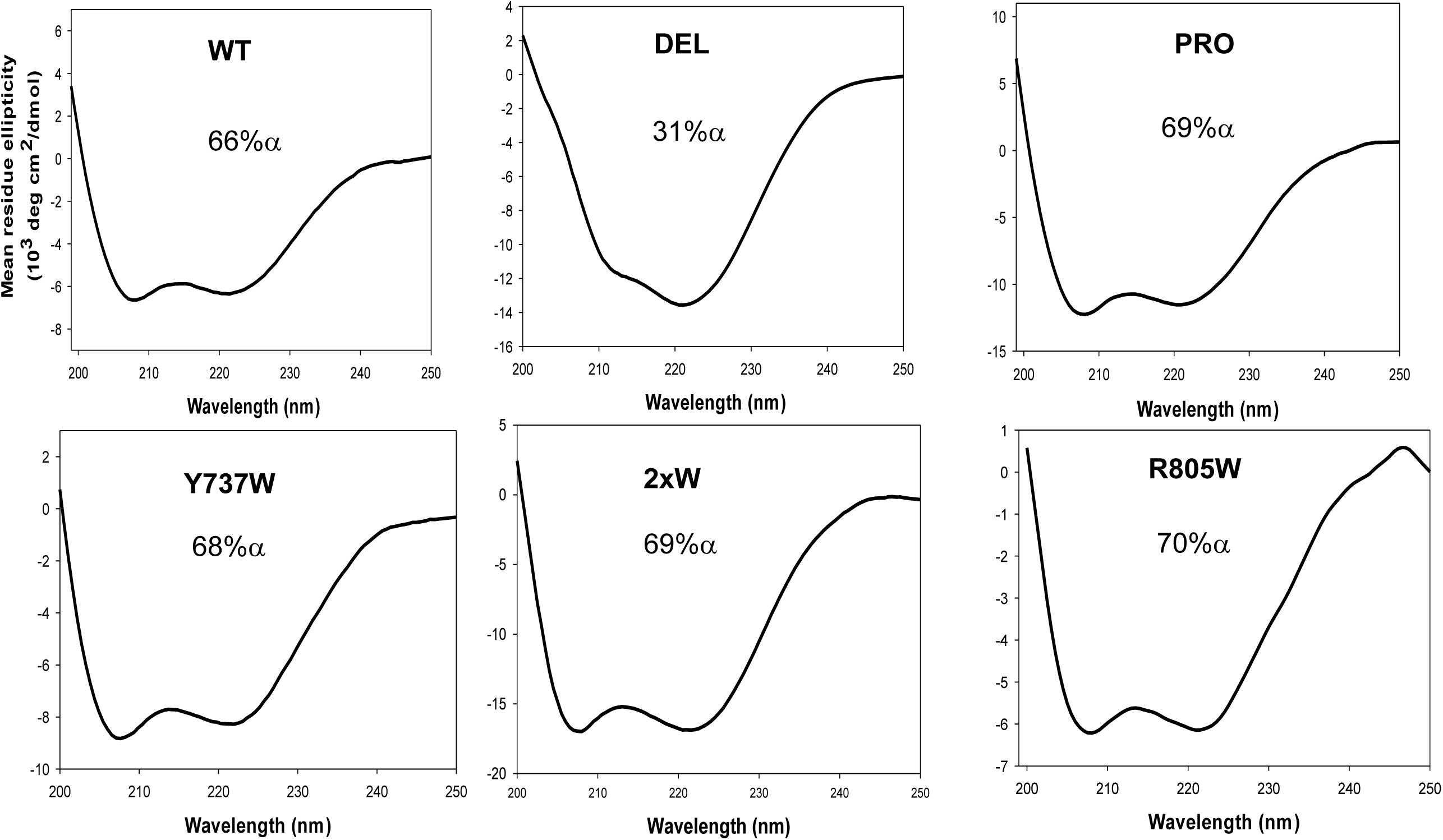
**Secondary structure of wild type and mutant UCS determined by circular dichroism.** The overall secondary structure determined by Far-UV circular dichroism of the WT, DEL, PRO, Y737W, 2xW and R805W constructs. The protein concentration was 1 μM in 30 mM phosphate buffer pH 7.4, 100 mM KCl, 1 mM MgCl_2_, 1mM TCEP buffer. A 0.1 cm path length cuvette was used. The % of alpha-helical content was estimated by using the K2D3 tool (63), and these are shown in each graph.

To assess the impact of the mutations on the overall stability of the UCS domain we measured the temperature dependence of the CD spectra. The thermal denaturation curve for the WT UCS domain measured at 222nm (**Fig. 5**) shows a cooperative transition with a Tm of 39 °C. The CD-spectra obtained at 60 °C shows that the secondary structure of all proteins is almost completely lost (and predominantly random coil), demonstrating that all had reached the same unfolded state (not shown). Cooling of the proteins back to 20 °C resulted in the original signal, indicating that thermal unfolding is fully reversible.

**Figure 5.**
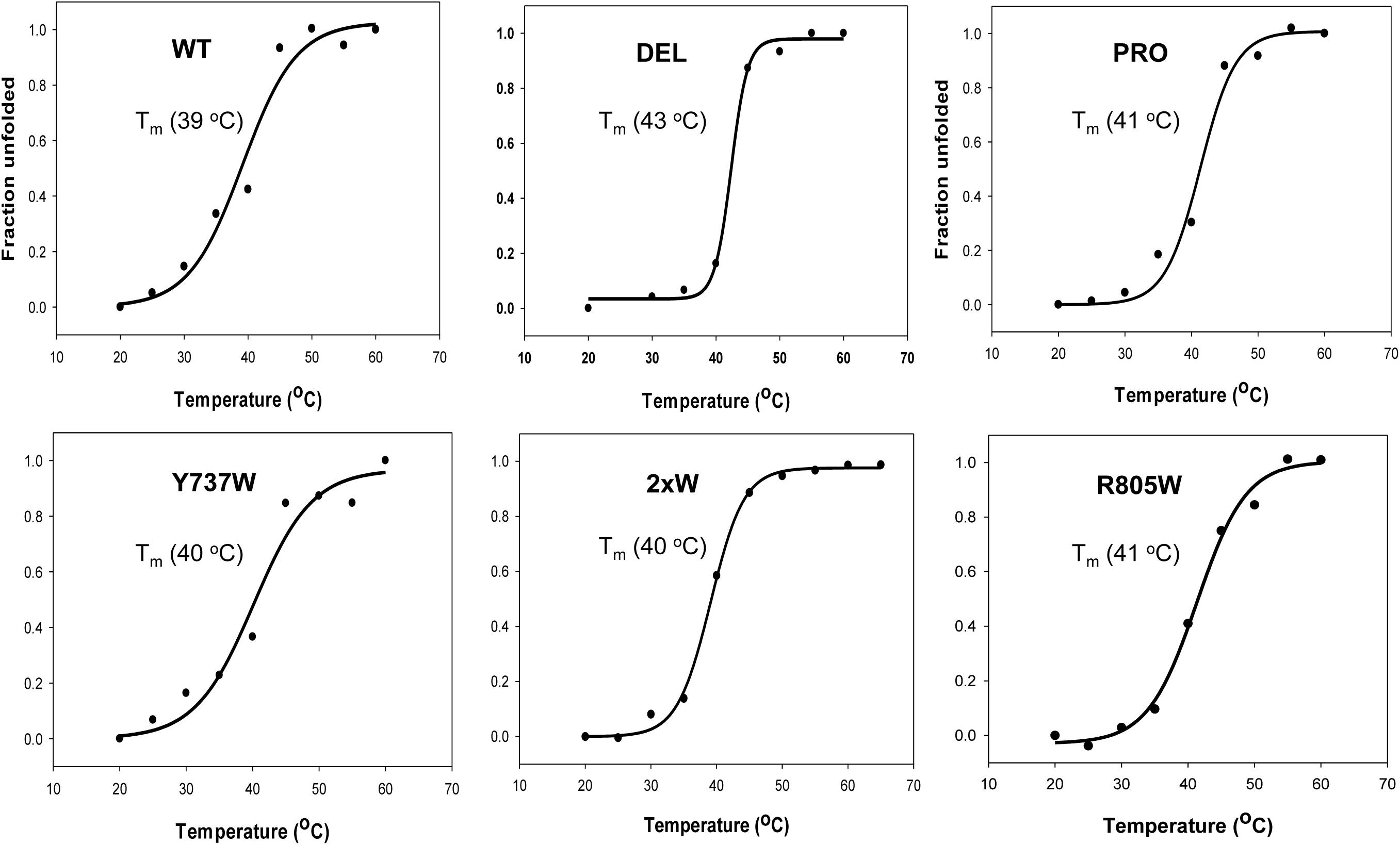
**Thermal denaturation of WT and mutant UCS proteins monitored by circular dichroism.** UCS proteins at 1uM were heated from 20 to 60 °C, and their denaturation was followed by circular dichroism ellipticity at 222 nm. The solid lines represent a fit to a 2-state denaturation model as described in the MATERIALS AND METHODS section.

The solid lines in **Fig. 5** show a fit to a simple 2-state unfolding cooperative model (26, 27). As seen in **Fig. 5** the curves for the four mutant proteins PRO, Y737W, 2xW and R805W are very similar to the WT with a Tm of ∼40°C. In contrast, the denaturation curve for the DEL mutant protein has a steeper transition with a Tm of ∼43°C, indicating a more cooperative unfolding transition than the WT protein (42). This result is consistent with the CD data (Fig. 4) and suggest the DEL mutant has a distinct structure, with an increase beta-sheet component.

### Temperature-induced conformational changes of WT and mutant UCS proteins monitored by Trp fluorescence

We further investigated temperature-induced structural changes in the UCS domain by tryptophan fluorescence. Steady-state Trp fluorescence spectroscopy is commonly used to track conformational transitions in proteins as a function of the temperature (43–45). The UCS domain has three Trp residues, W517, W530 and W847, thus enabling tracking the global conformational changes of the UCS domain. A surface representation of the three Trp residues reveals that only one residue (W847) is partially buried in the structure (**Fig. 6A, top diagram**). We found that the relative Trp fluorescence signal for the WT UCS shows a gradual decrease as the temperature was raised, with a Tm of 27°C (**Fig. 6A**) indicating a change from a hydrophobic environment to one of aqueous solvent. Most of this signal likely originates from W847 since the other two Trp residues are solvent exposed. The melting curve for PRO was not significantly different from WT and had a Tm of 27 °C. In contrast, the melting curve for DEL had a significant shift of the Tm by 9°C, indicating a stabilization of the overall structure as reported by residue W847.

**Figure 6.**
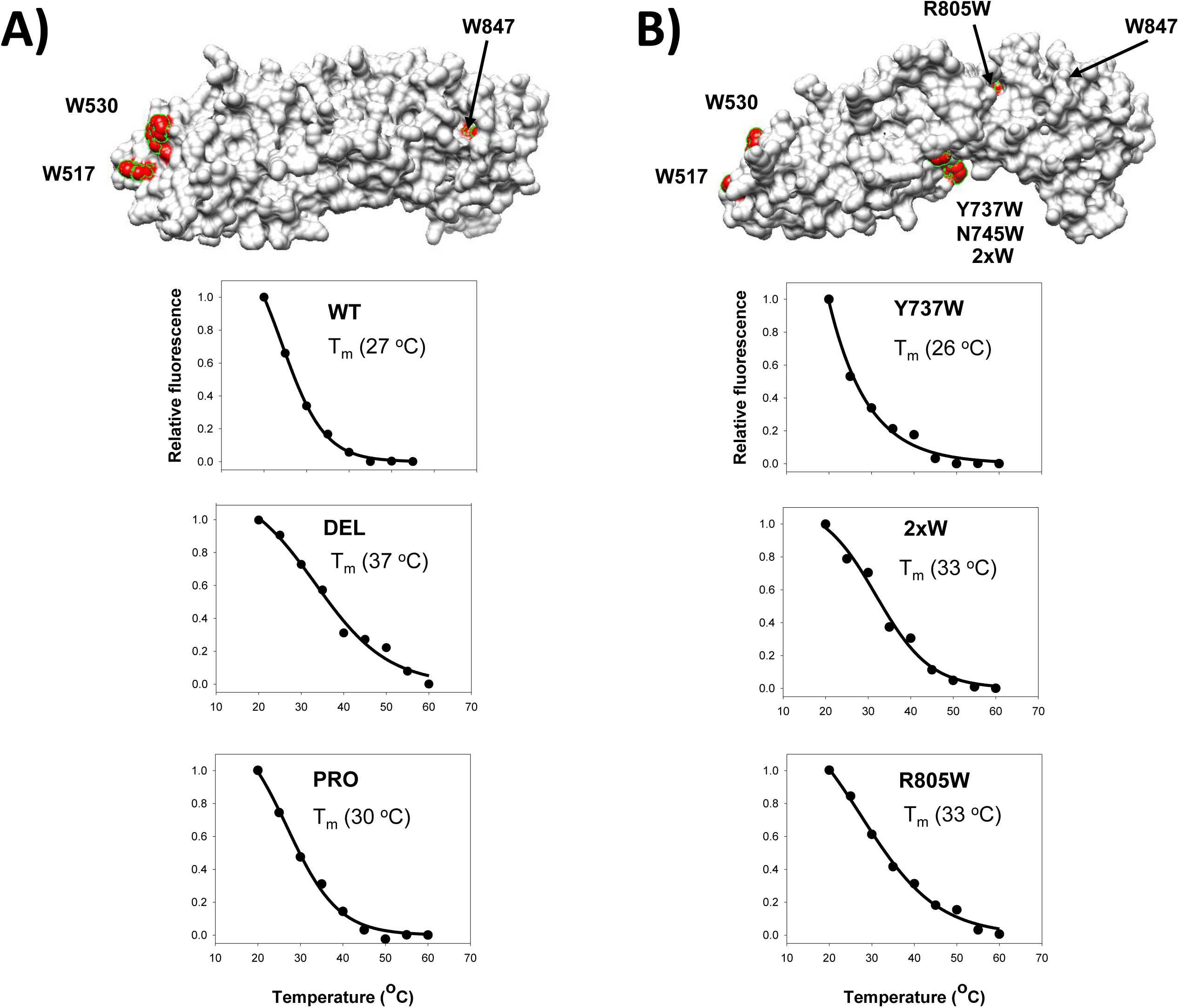
**Temperature-induced conformational changes of WT and mutant UCS proteins monitored by Trp fluorescence.** UCS proteins at 1uM were heated from 3 to 89 °C, and their denaturation was followed by fluorescence at 330 nm with excitation at 280 nm. **A)** The diagram shows a surface representation of the UCS domain highlighting the three Trp residues. The plots show the temperature curves for the WT, DEL and PRO proteins. **B)** The diagram shows a surface representation of the UCS domain highlighting the three Trp residues plus the engineered Trp mutations. The plots show the thermal curves for the Y737W, 2xWW and R805W proteins.

**Figure 6B** shows the temperature-induced conformational changes of the mutant UCS proteins with extra Trp residues (Y737W, 2xW, R805W). A surface representation of the extra Trp residues reveals that only one extra residue (R805W) is partially buried in the structure (**Fig. 6B, top diagram**). So, most of the Trp signal likely originates from the W847 and W805 residues. However, we cannot exclude contributions from residues Y737W and N745W. The temperature-dependence curve for Y737W was very similar in shape to that of WT and had a Tm of 26°C. The curve of the double Trp mutant (2xW) had a Tm of 33°C, suggesting a stabilization of the UCS structure by the N745W mutations with respect to the Y737W; the temperature-dependence curve for R805W is similar to 2xWW with a Tm of 33°C.

These data suggests that the DEL mutation increases the overall structural stability of the UCS domain, suggesting that this mutant has a distinct conformation.

### Measuring chaperone activity of mutant UCS proteins using a S1 Mg-ATPase heat inactivation protection assay

To evaluate the effect of mutations on the chaperone activity of the UCS mutants, we measured how effective these proteins would be in protecting S1 domain unfolding (or partial unfolding) when exposed to heat stress. We assessed unfolding under heat-shock conditions by determining the effect of elevated temperatures on its Mg-activated ATPase activity. Residual Mg-ATPase activity of S1 protected by the UCS chaperones were compared to the S1 alone and the relative activity is shown in **Fig. 7**, where S1 Mg-ATPase activity after 10 minutes at 42°C is set as 100%. As the data show, the full length UNC-45B and the UCS domain had the strongest heat protection effect (∼2200% and ∼2600%, respectively) whereas all mutants had a significantly lower heat protection effect on ATPase activity (DEL: ∼1000%, WW: ∼1600%). These data support our hypothesis that the client-holding loop and conserved residues in the client-binding groove play an important role in the UCS domain chaperoning activity. Interestingly the R805W mutation, which is far away from the client-binding groove, also show a relatively weak chaperoning activity compared to WT.

**Figure 7.**
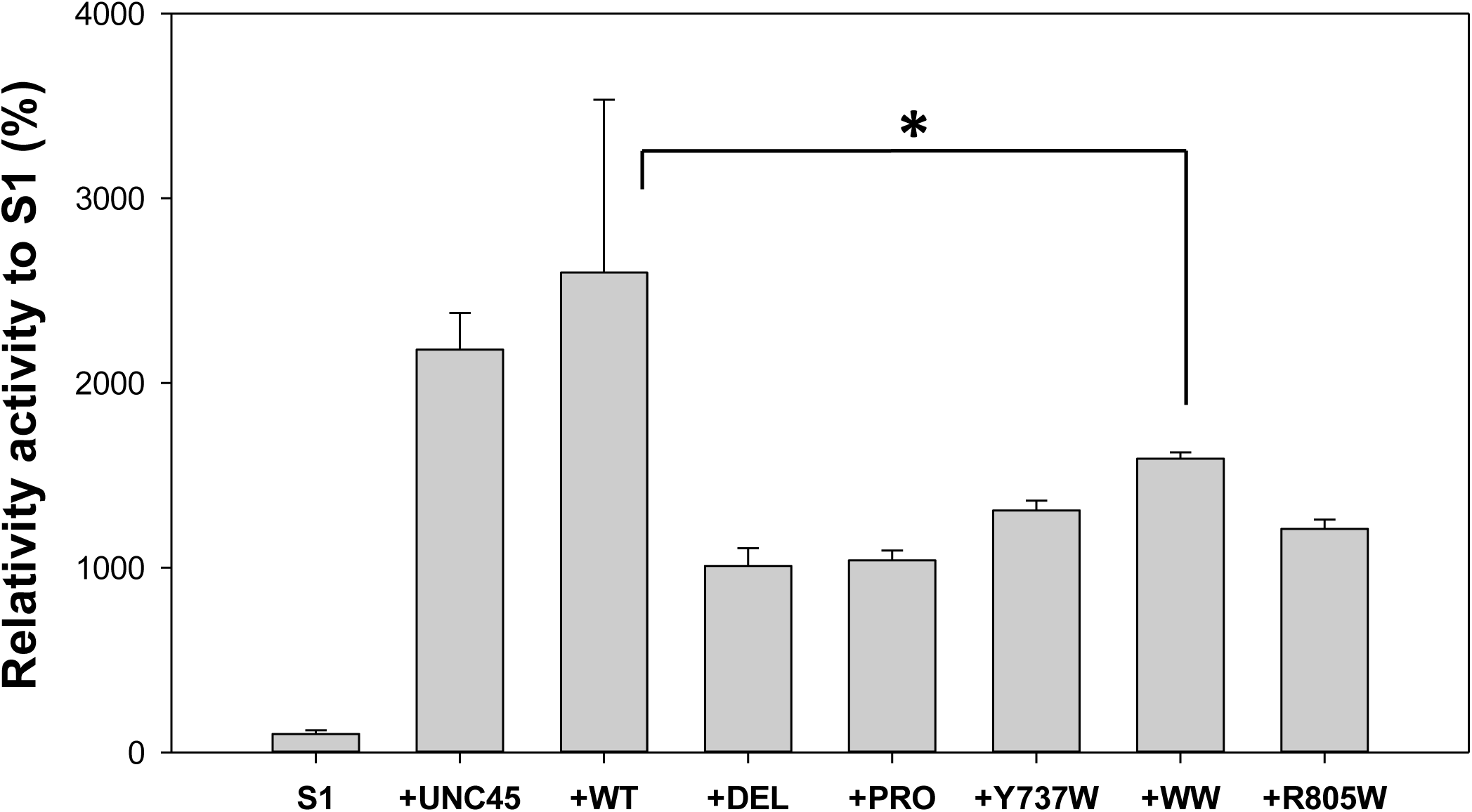
**Protection of S1 Mg-ATPase activity from heat by UNC45B, UCS wild-type and the UCS mutants**. UNC45B, UCS and its mutant variants were incubated in five-time excess with S1 at 42°C for 10 min. Residual Mg-ATPase activity of S1 protected by the chaperones were compared to the S1 alone and relative activity is displayed in the graph, where S1 Mg-ATPase activity after 10 minutes at 42°C is set as 100%.

### Measuring chaperone activity of mutant UCS proteins using a heat inactivation Citrate Synthase assay

To further test the effect of mutations on the chaperoning activity of the UCS domain, we used citrate synthase (CS) as a chaperone substrate (33, 46). CS has been extensively used as a model substrate for the four major classes of molecular chaperones (small heat shock proteins (Hsps), GroEL, Hsp70, and Hsp90 (33, 46). UNC-45 also prevents heat-induced aggregation of citrate synthase and facilitates its refolding (11, 47) suggesting that like other ATP-independent chaperones (48), UNC-45 can interact with several client proteins. CS is very sensitive to thermal inactivation; its activity is almost completely lost following incubation at 42°C for 10 min and the inactivation of CS followed apparent first-order kinetics (**Fig. 8A, black circles**). The presence of UNC-45B during the incubation of CS at 42°C changed the inactivation kinetics significantly (**Fig. 8A, Grey triangles**). In the presence of a 10-fold molar excess of UNC-45B the obtained apparent rate constant of inactivation decreased by 4-fold (from 0.2 min^-1^ to 0.05 min^-1^). Control experiments using 10:1 BSA/CS ratio showed that BSA also protects but at much lower extent (about 120% of relative CS activity; data not shown).

**Figure 8.**
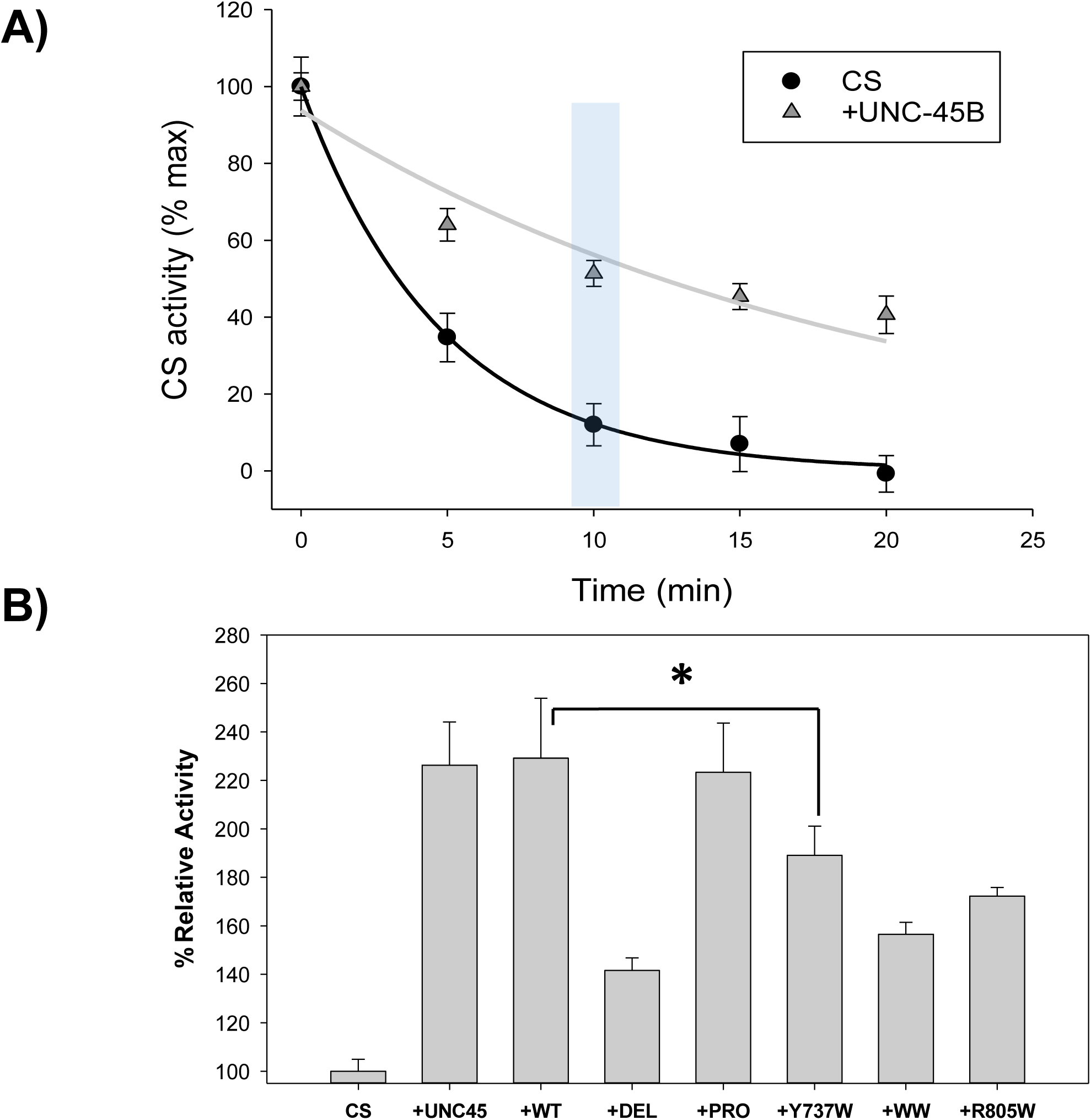
**Influence of wild type and mutant UCS proteins on the thermal inactivation of citrate synthase.** UNC45B, UCS and its mutant variants were mixed with CS and diluted in 10:1 ratio in HEPES-KOH, pH 7.4 or 20 mM Tris HCl, 100mM KCl, pH 8.0 (at a concentration of 2 µM and 0.2 µM, respectively). Each mix, of CS and chaperones, as well sample of CS alone were incubated for 10 min at 42°C. Upon heat shock, samples were cooled down to 4°C, briefly centrifuged and CS activity was performed. Assays were run in triplicates in at least two independent separate experiments.

Using this chaperone assay, we tested our UCS mutants by measuring the relative CS activity after incubation for10 min at 42°C (marked by the cyan bar in **Fig. 8A**). As the data shows the DEL mutant has the weakest protection effect of CS from thermal inactivation (3.3-fold lower than WT). The mutations that target the client binding groove, Y737W and 2xW, had a significantly lower protecting effect than WT. Unexpectedly, and in contrast with the S1 assay, the holding loop PRO mutant was very similar to WT in protecting CS activity from heat. This suggest that the “trap door” function of the UCS holding loop may depend on particular residues of the exposed client peptides within the groove (more specific for S1 myosin than CS). These data further support our hypothesis that conserved residues in the client-binding groove play a key role in the UCS domain chaperoning activity. Interestingly, as with the CS assay the R805W mutant also displayed a relatively weak chaperoning activity compared to the WT UCS.

### Deletion of the client-holding loop leads to a large change in shape of the UCS domain

The CD and intrinsic fluorescence data clearly show that the DEL mutant has a different secondary structure and higher structural stability compared to the WT UCS protein **(Figs. 4 and 5**). Also, the thermal protection data of the activity of both client proteins, myosin-S1 and CS, show that the DEL mutant has a weaker chaperone activity than the WT protein (**Figs. 7 and 8B**). In order to better understand these findings, we employed small angle X-ray scattering to determine the solution conformations of the DEL and WT proteins.

**Figure 9A and 9B** show *ab initio* molecular shapes of the WT and DEL UCS proteins, respectively. The best models were obtained by assuming a dimer conformation. The WT protein appear to dimerize through interactions between loops at the end of helices H2-H3 of the first armadillo repeat in the UCS domain via a salt-bridge (**Fig. S1D**). The dimer interface in the DEL dimer has moved from the first armadillo repeat and is larger, spanning two ARM repeats. A dilution series displays a concentration dependence consistent with a monomer-dimer equilibrium for the WT UCS (**Fig. 9C**, red squares and line), with an estimated Kd= 0.53mg/ml. In contrast, the dilution series for the DEL mutant is consistent with a dimer at each concentration (**Fig. 9C**, blue diamonds and line), with an estimated Kd < 0.01 mg/ml. In order to test the hypothesis that the WT protein dimerize via electrostatic interactions (**Fig. S1D**), we increased the salt concentration to suppress ionic interactions. Indeed as **Fig. S3** shows, the fraction of dimer in solution dramatically decreases when adding 1M LiCl.

**Figure 9.**
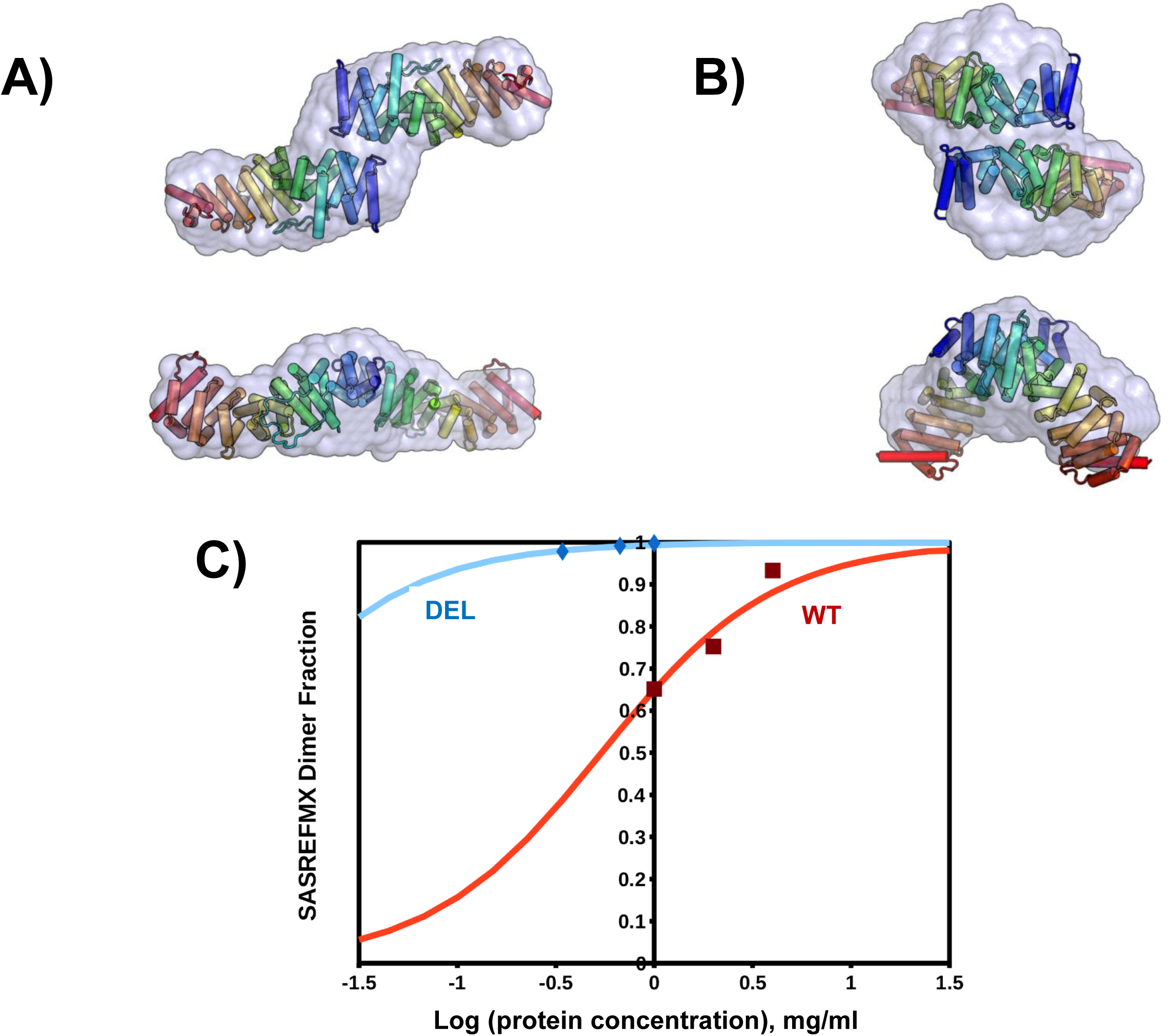
**SAXS Models of the WT and DEL proteins. A)** The WT SASREFMX fit (X2=1.1,1.0,1.0) and mutual dimer shown with the 4 mg/ml Gasbor model surface. **B)** The DEL 0.5 mg/ml merged data CORAL fit (χ2=1.1) and dimer model, with Gasbor (surface). The two images show the same structure rotated by 90 degrees. **C)** Semi-Log Dimer fraction curves: The WT SAXS data (red line) (Kd=0.53 mg/ml) and DEL SAXS data (blue line) (Kd<<0.01 mg/ml).

The SAXS data suggest that deletion of the loop leads to a significant conformational change of the UCS domain and the formation of a stable dimer. This large change may hinder the access of the unfolded or partially folded client protein to the groove.

### Dominant-negative inhibition of thick filament assembly by expression of mutant UCS proteins in *C. elegans*

We wished to determine the *in vivo* effects of the expression of UNC-45 carrying these mutations in the UCS domain. In *C. elegans*, null mutations in the *unc-45* gene are “Pat” embryonic lethal, whereas amino acid substitutions in the UCS domain result in adults that show decreased accumulation of myosin heavy chains, decreased and disorganized thick filament assembly (49–51). Using the alignment of human and *C. elegans* UNC-45 UCS domains (**Fig. 2B**), we identified residues in the nematode sequence that are comparable to the residues mutated in the human UNC-45B UCS domain that we are studying. We attempted to use CRISPR/Cas9 gene editing to generate several of these mutants in the *C. elegans* genome, without success; in several cases we could identify heterozygotes, but could not isolate homozygotes, probably because of embryonic lethality. Therefore, we undertook a different strategy in which we utilized transgenic worms to assess the dominant negative *in vivo* effects of four different UCS domain mutations – Groove (LTNL→ AAVQ), R805W, Pro (QFAKH→ PPGPP), and 2xW (Y750W,N758W). We generated nematodes carrying extrachromosomal arrays with either wildtype or mutated HA tagged *unc-45* under the control of the body wall muscle specific *myo-3* promoter and used *sur-5::GFP* as the transgenic marker. Three independently generated transgenic lines were studied for each mutant protein. Animals were grown to adults, fixed, and immunostained with antibodies to myosin (MHC A; red in **Fig.10**), to GFP, and to UNC-95 (both green in **Fig. 10**). Because such transgenic lines show somatic mosaicism, i.e. not every muscle cell receives the array and expresses the protein, we identified which muscle cells actually express the protein, by finding those that also expressed the transformation marker fused to GFP (cells with green nuclei in **Fig. 10**). In addition, to locate the region of the muscle cell that normally contains the thick filaments, we immunostained with anti-UNC-95, which localizes to the base of integrin adhesion complexes in muscle (also green in **Fig. 10**). All three lines expressing wt UNC-45 displayed normal thick filament assembly and organization with parallel A-bands, indicating that the extrachromosomal *unc-45* was not being overexpressed to the point of causing a phenotype. Two out of the three Groove and Pro mutation lines created displayed disorganized thick filaments. All three R805W and 2xW lines created displayed disorganized thick filaments in the muscle cells expressing the extrachromosomal array. The R805W mutation, which disrupts a salt bridge between helices 15H2 and 14H2, appeared to have the most drastic effect on the structure and organization of the worm sarcomeres. The mutant protein expressed in the two lines (Groove line 2 and Pro line 2) that showed relatively normal thick filament organization may not have been expressed at a high enough level to give a dominant phenotype or the mutant protein was so unstable that it was degraded quickly after translation. We used an antibody to the HA tag, along with antibodies to Myosin A and GFP, to verify expression and localization of the HA-tagged UNC-45 protein in the different strains (**Supplemental Figure S4**). Wherever the thick filament structure and organization remained intact, the HA antibody can be seen localizing as striations to either side of Myosin A, which is normal UNC-45 localization. However, in the mutants wherever the thick filaments were disrupted, the HA-UNC-45 failed to localize properly and appeared to either be aggregating or reduced compared to the intact portions of the muscle cell. This result suggests that the mutant UNC-45 proteins act as dominant negative poisons. Perhaps this occurs by binding to the normal endogenous UNC-45, and reducing the overall concentration of functional UNC-45, so that myosin heads are not properly folded, and thick filament assembly and organization are affected. This would be consistent with the fact that in crystal structures, UNC-45 has been found in oligomers (17, 20). These results provide evidence for the biological importance of these UCS regions/residues within the UNC-45 chaperone.

**Figure 10.**
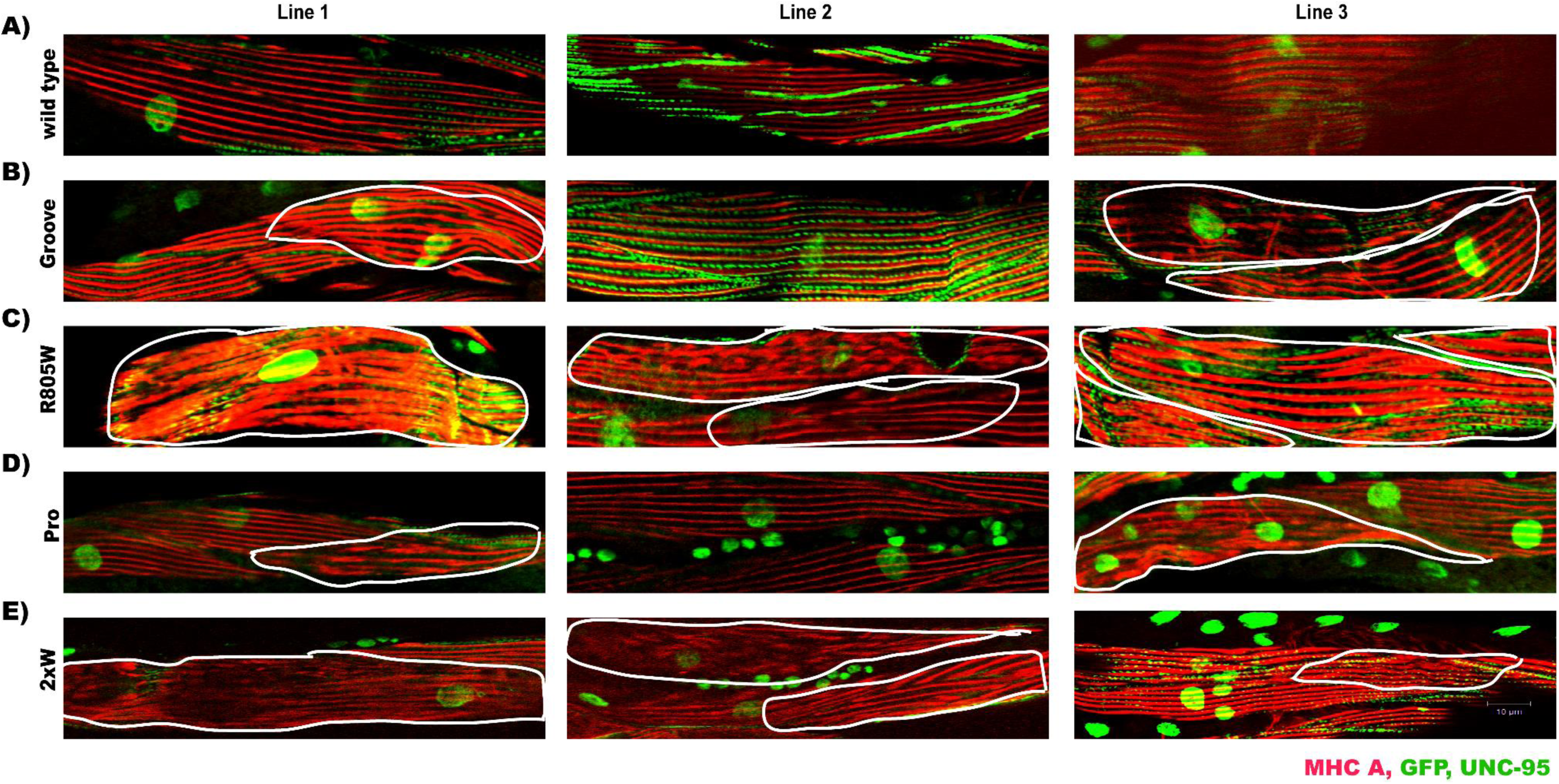
**The effects of expression of UNC-45 containing mutations in the UCS domain *in vivo*.** Immunostaining of three *C. elegans* lines expressing either extrachromosomal Wildtype (A), Groove (B), R805W (C), Pro (D), or 2xW (E) UNC-45. Myosin A (MHC A), which localizes to the A-bands of the muscle cells, is shown in red. UNC-95, which localizes to the base of the integrin adhesion complexes in muscle cells and was used to identify the myofibrillar region of muscle cells independent of A-band organization, is shown in green. GFP, which localizes to the nucleus of all muscle cells expressing the transgenic array (transformation marker *sur-5*::GFP), is also shown in green. White boundaries have been drawn around areas of A-band disruption/disorganization.

## 4. DISCUSSION AND CONCLUSIONS

Here we used a combination of biophysical and biological experimental tools to obtain a better understanding of the molecular mechanisms by which the UCS domain of UNC-45B chaperones its client. We engineered mutations in the chaperone domain of UNC-45B (listed in Table I) designed to: i) disrupt chaperone-client interactions by removing and altering the structure of the putative client-interacting loop (DEL and PRO mutants, respectively) and ii) disrupt chaperone-client interactions by changing highly conserved residues in the putative client-binding groove (Groove, Y737W, N745Y mutants). We also introduced the R805W mutation to test the hypothesis that the stability of the salt bridge between R805-E767 plays a role in UCS-client interactions (25).

Our biophysical data show that removing the holding loop (DEL mutation) had the most pronounced effect on the secondary structure and thermal stability of the UCS domain **(Figs. 4 and 5**); there is a significant decrease in alpha helix character (from 68% to 31%) and increase in thermal stability (a Tm shift from 39° to 43°). Also, the DEL mutant has the weakest protection effect on CS from thermal inactivation (3.3-fold lower than WT, **Fig. 8**). The SAXS data revealed that deletion of the loop leads to a significant conformational change of the UCS domain and the formation of a stable dimer (**Fig. 9**). This large change may hinder the access of the unfolded or partially folded client protein to the groove. These results are consistent with the *in vivo* findings by Gazda et. al. (17) that deletion of this loop mutation neither rescue the defect in *C. elegans* sarcomere organization nor bind to myosin in immunoprecipitation experiments.

Our *in vivo* data show that, to our surprise, the R805W mutation appeared to have the most drastic effect on the structure and organization of the worm sarcomeres (**Fig. 10** and **Fig. S4**). This mutation does not affect UCS secondary structure (**Fig. 4**) or structural stability (**Figs. 5 and 6**) but significantly impairs its chaperoning activity (**Figs. 7 and 8**). The facts that the autosomal dominant mutation R805W segregates with human congenital/infantile cataract and that the R805W mutant form of human UNC45B expressed in zebrafish embryonic eye results in aberrant lens maturation suggest a crucial role of R805 in UCS domain stability and/or client interaction, which in the case of developing lens is non-muscle myosin (25); based on the x-ray structure of Drosophila UNC45, R803 (Hs R805) has been implicated to form inter-repeat ion pair interaction with Dm E766 (Hs E767). Consistently, *C. elegans unc-45(m94)* temperature sensitive mutants with affected locomotion and sarcomere integrity has missense mutation in E781, which is one of the E780/781, located in the corresponding area to human E767. *C. elegans* has two glutamines in positions 780 and 781, while other species including Drosophila, zebrafish, mouse and human have only one glutamine. Interestingly, one of the first identified *C. elegans* ts mutations in UNC45, which affects worm motility and sarcomere organization, is a point mutation in the corresponding codon resulting in E781K substitution (52). It has been recently found that that the worm UNC45 E781K mutant protein has different unfolding Tm as measured by CD, compared to other UCS ts mutants (20), suggesting that this mutation affects the structure of the UCS domain

The Y750W (hs Y737W) mutation itself doesn’t drastically affect chaperoning ability of UNC45 as shown in rescue experiments with worms (17) and myosin motor domain folding assay in insect cells (20). Our data show that the double mutant 2xW (Y737W, N745W) significantly affects the *in vitro* chaperoning function of UNC-45 (**Figs. 7 and 8**) and also greatly disrupted thick filament organization in worms (**Figs. 10 and S4**).

Our results may have offer insights into to the folding and repair of the many classes of myosin molecules that are expressed in non-muscle tissues. In fact, the expression and function of UNC-45 is not limited to just striated muscle, but is very likely to be crucial for the folding of myosin heads of the large myosin superfamily containing at least 35 different classes of myosins, some of which are expressed in nearly all cell types. UNC-45 is required for early embryonic development in *C. elegans* (53, 54), and is involved in cell division and cell migration of ovarian and breast carcinoma (epithelial cells) (55–57); in both of these cases, UNC-45 likely aids the folding of non-muscle myosin II. A better understanding of the mechanisms by which UNC-45 helps fold myosin heads is likely to lead to new therapies for a number of diseases, especially those due to mutations in myosin.

## Author Contributions

A.F.O. and G.M.B. conceived the study; I.V. and T.M. expressed and purified proteins; M.V., S.P., I.V., T.M. performed CD and fluorescence experiments and analyzed data with A.F.O.; I.V. T.M. performed chaperone experiments and analyzed data with A.F.O.; M.W. performed SAXS experiments and analyzed data; H.Q., C.J.C. performed worm immunofluorescence experiments; I.G., G.M.B. and A.F.O., wrote the manuscript.

## ACKNOWLEDGMENTS

This work was supported by National Institutes of Health grant R01GM118534 to G.M.B. and A.F.O.

